# iLID-antiGFP-nanobody is a flexible targeting strategy for recruitment to GFP-tagged proteins

**DOI:** 10.1101/2022.11.24.517828

**Authors:** Eike K. Mahlandt, Maarten Toereppel, Tayeba Haydary, Joachim Goedhart

## Abstract

Optogenetics is a fast-growing field, that applies light-sensitive proteins to manipulate cellular processes. A popular optogenetics tool is the improved light-induced dimer (iLID). It comprises two components, iLID and SspB, which heterodimerize upon illumination with blue light. This system is often used to recruit proteins to a specific subcellular location, e.g. by targeting the iLID to the plasma membrane. The targeting requires modification of the iLID with a targeting sequence. To skip the modification of the iLID and use existing GFP fusion as targets, we fuse an antiGFP nanobody to the iLID. We show that the antiGFP nanobody is able to locate iLID to GFP-tagged proteins. Plus, the light-dependent recruitment of SspB to iLID, localized by the antiGFP nanobody to a GFP-tagged protein, is still functioning efficiently. This approach increases flexibility, enabling the recruitment of any GFP-tagged protein, without the necessity of protein engineering.

## Introduction

Optogenetics uses light-sensitive proteins to manipulate cellular processes, inducible by light. This provides the advantages of global to subcellular induction, reversibility of this induction and non-invasive activation with light. One well-established optogenetic tool is the improved light-induced dimer (iLID), a heterodimerization tool (Guntas et al., 2015). It consists of the binding pair SspB and SsrA, where SsrA is shielded by the LOV domain (together called iLID) in the dark state. Light induces a conformational change in the LOV domain, followed by the release of the SsrA domain. This domain becomes now available for binding to SspB and based on diffusion, the SspB binds SsrA. This effect can be applied to recruit a protein of interest (POI) to the location of iLID, in a local and reversible manner. This system has been used to manipulate signaling pathways. For example, the catalytic active domain of Rho guanine-nucleotide exchange factors (GEF) has been recruited to the plasma membrane to induce Rho GTPase signaling and thereby alter the cell morphology, e.g. the formation of local lamellipodia (Guntas et al., 2015).

These experiments require overexpression of the signaling molecules fused to SspB but also the target protein, which by itself might already have a signaling effect. iLID is widely used and often in combination with a plasma membrane tag (Dessauges et al., 2022; Gonzalez-Martinez et al., 2022; Saha et al., 2022; Vaidžiulytė et al., 2022). The approach can be improved by targeting proteins to a more specific location, for example recruiting to focal adhesions or adherens junctions where local GEF and GAP activity have been detected (Müller et al., 2020). However, the protein engineering and overexpression of the relevant molecules have been proven challenging and difficult.

A solution to such issues can be provided by nanobodies, small antibodies (12-15 kDa) that can specifically recognize proteins. These nanobodies provide a way to tag endogenous proteins with the iLID system to manipulate the location and/or signaling of endogenous proteins. A well-characterized nanobody is the antiGFP nanobody, originally called nanotrap with a size of 13 kDa (Rothbauer et al., 2008). Rothbauer and colleagues also showed that the GFP-tagged proteins can be targeted to a nuclear envelop localized antiGFP nanobody. Other studies have applied the antiGFP nanobody to link a GFP-tagged protein to a quantum dot (Derivery et al., 2017) or APEX tag for electron microscopy (Ariotti et al., 2015) or additionally to alter the location of a GFP-tagged protein (Derivery et al., 2015; Schornack et al., 2009). More recently, an entire toolkit of functionalized nanobodies against GFP and RFP has been developed to visualize and manipulate cell signaling (Prole and Taylor, 2019). This study includes the nanobody-mediated targeting of the optogenetic dissociation tool LOV2/zdk1 to mitochondria. Furthermore, the antiGFP nanobody has been used in the optogentic CRY2/CIBN based light-activated reversible inhibition by assembled trap (LARIAT) that inactivates proteins by clustering them in large complexes (Osswald et al., 2019; Park et al., 2017). To the best of our knowledge, the iLID system has not been used in combination with an antiGFP nanobody.

We propose any GFP-tagged protein could be used in combination with the antiGFP nanobody iLID system, to save time and effort in protein engineering or generating stable cell lines and providing more variety in iLID targets. Here we present a proof-of-principle experiment showing that the antiGFP nanobody can be applied to target iLID to a GFP-tagged protein and efficiently and specifically recruit SspB.

## Results

To evaluate if the antiGFP nanobody can be applied to locate iLID to a GFP tagged protein, we tested which fluorescent proteins are bound most efficiently by the nanobody. From here on we will refer to the antiGFP nanobody as nanobody. The nanobody was tagged with a fluorescent protein that was not derived from avGFP, e.g. mScarlet or mNeonGreen, and as we expected it had hardly or no affinity for these fluorescent proteins. To evaluate binding, we used a target fluorescent protein that was fused to histone 2A to localize it in the nucleus. The nanobody and target were co-transfected and we measured co-localization in individual cells. The nanobody strongly co-localized with EGFP and LSS-sGFP2 (**Fig. 1A,B**). The nanobody fused to the N-terminus of mScarlet-I seems to have a binding advantage to EGFP over the nanobody fused to the C-terminus of mScarlet-I. mTurquoise2, developed from the avGFP ancestor, is also bound by a small fraction of the nanobody (**Fig. 1A,B**). Fluorescent proteins, developed from other precursors than avGFP, such as mCherry, mScarlet-I, iRFP670, mKate2 and the green fluorescent protein mNeonGreen were not bound by the nanobody (**Fig. 1A,B**). Therefore, these proteins can be used as fluorescent labels in combination with the nanobody. Moreover, avGFP-related proteins can serve as a target for the nanobody.

**Fig. 1.**
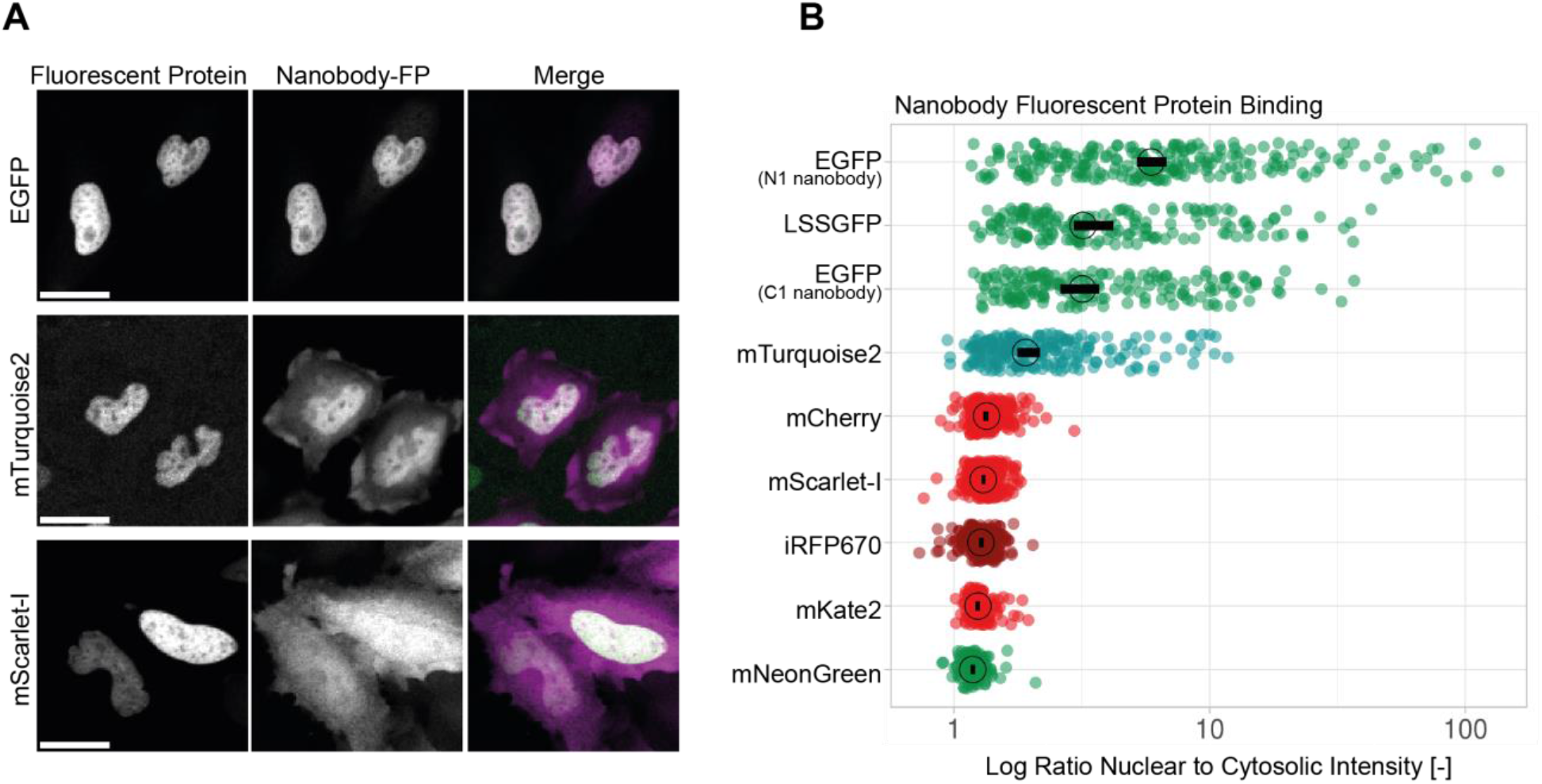
Nanobody specificity for fluorescent protein binding. (**A**) Confocal images of HeLa cells co-expressing EGFP-H2A + nanobody-mScarlet-I-N1 (top), mTurquoise2-H2A + mScarlet-I-nanobody-C1 (middle) and mScarlet-I-H2A + mNeonGreen-nanobody-C1 (bottom), showing the nuclear tagged fluorescent protein on the left, the fluorescently tagged nanobody in the middle and a merge on the right side, where the single of the fluorescent protein is shown in green and of the nanobody in magenta. Scale bars: 25 μm. (**B**) Binding of the AntiGFP-nanobody to different fluorescent proteins measured as the ratio of nuclear to cytosolic intensity for HeLa cells co-expressing EGFP-H2A + nanobody-mScarlet-I-N1, H2A-LSSsGFP2 + mScarlet-I-nanobody-C1, EGFP-H2A + mScarlet-I-nanobody-C1, mTurquoise2-H2A + mScarlet-I-nanobody-C1, mCherry-H2A + mNeonGreen-nanobody-C1, mScarlet-I-H2A + mNeonGreen-nanobody-C1, iRFP670-H2A + mNeonGreen-nanobody, mKate2-H2A + mNeonGreen-nanobody and mNeonGreen-H2A + mScarlet-I-nanobody. Each dot represents the measurement of one cell. The median of the data is indicated as a circle. The 95% confidence interval is indicated by a black bar. The number of cells measured in two experiments based on independent transfections is: EGFP-(N1 nanobody)=224, LSSGFP=164, EGFP (C1 nanobody)=168, mTurquoise2=206, mCherry=202, mScarlet-I=234, iRFP670=152, mKate2=116, mNeonGreen=104.

As a proof-of-principle experiment to show that the nanobody fused to iLID functions as a linker between a GFP-tagged protein and the recruitment of SspB, HeLa cell were transfected with either the nuclear tag EGFP-H2A, the endoplasmic reticulum (ER) tag CytERM-mEGFP or the plasma membrane tag EGFP-CaaX and the (antiGFP)-nanobody-HaloTag-iLID and SspB-mScarlet-I. The (antiGFP)-nanobody-HaloTag-iLID construct localized at the same location as the GFP-tagged markers. Upon photo-activation, the SspB-mScarlet-I was recruited to these locations (**Fig. 2A,B,C**). Interestingly, the recruitment seemed to plateau after roughly 30 s for the ER and plasma membrane set up, where for the nuclear set up it showed continues recruitment until the end of the recording, except for the mitotic cell. This delay might be explained by the limited transport via the nuclear envelope, which was broken down in the mitotic cells, therefore allowing free diffusion of SspB to the location of iLID.

**Fig. 2.**
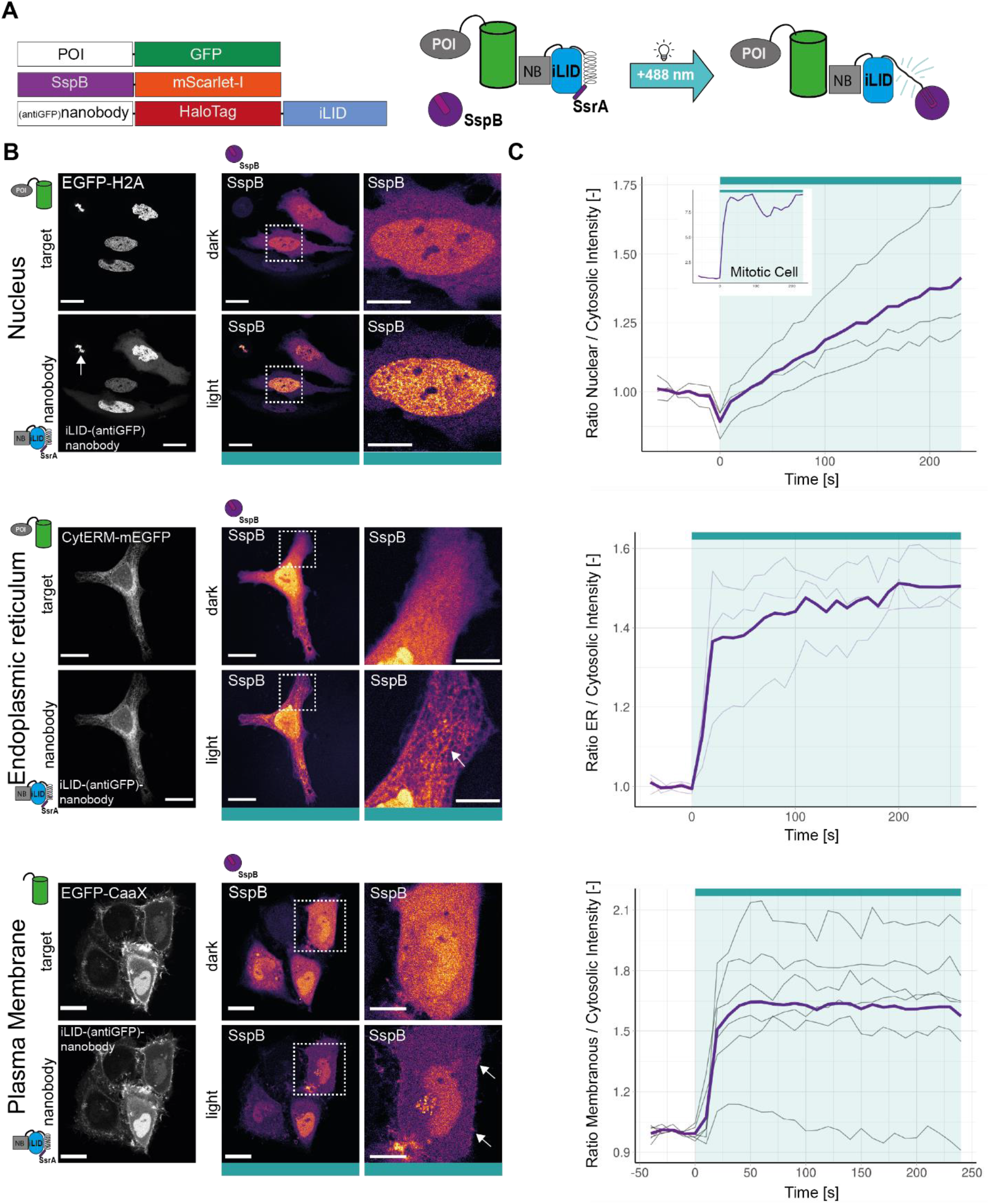
SspB recruitment to different GFP tagged targets via iLID-nanobody. (Legend on the following page) (**A**) Schematic of the three constructs and their interactions. POI=protein of interest, NB=nanobody. (**B**) Confocal images of HeLa cells co expressing antiGFP-nanobody-HaloTag-iLID (grey) and SspB-mScarlet-I (LUT=mpl inferno) in combination with either EGFP-H2A (top), CytERM-mEGFP (middle) or EGFP-CaaX (bottom). SspB-mScarlet-I is shown before and after blue light photo activation indicated with cyan bar. Dashed box indicates zoom-in on the right side. First arrow indicates mitotic cell; the other arrows indicate SspB recruitment to the target structure. Scale bars: whole field of view 25 μm, zoom-in 10 μm. (**C**) Time traces of the ratio of target over cytosolic SspB-mScarlet-I intensity for the in **B** described cells. Thin black lines represent single cells. The purple line represents the mean. The cyan bar indicates photo activation by blue light.

This experiment showed that it is possible to use the nanobody to localize iLID at a GFP tagged protein and subsequently recruit SspB to this location induced by photo activation.

## Discussion

One of the most frequently used optogenetic heterodimerization systems is iLID. iLID is often targeted to the plasma membrane. To localize iLID to other targets, e.g., proteins or organelles, requires additional protein engineering and overexpression of such target proteins. One of the major drawbacks is that engineering of some proteins may be cumbersome and difficult. However, GFP fusions are available for many proteins, and for some there are cell lines stably expressing GFP-tagged proteins, of which some are expressed on the endogenous level (de Man et al., 2021). Here we show that GFP-tagged proteins can be utilized to locate an antiGFP-nanobody-iLID construct to a GFP-tagged protein and recruit SspB to this target location. This enables the targeting of iLID to GFP-tagged proteins without additional genetic engineering.

Here we demonstrated binding of the nanobody to EGFP and LSS-sGFP2. In general, if the fluorescent protein is similar enough to avGFP it can be used as target. A previous study confirmed using fluorescent microscopy that sGFP, mEmerald, Clover, and also EYFP, mVenus and mCitrine are bound by the antiGFP nanobody (Küey et al., 2019). Küey and colleagues also confirmed that RFPs did not bind the nanobody as was confirmed by us. However, they did find no binding of mTurquoise2, whereas we did detect binding of the nanobody but less efficient than to GFP. For an experimental setup, one may consider a target protein tagged with GFP or YFP. SspB and iLID can then be tagged with red and far red fluorescent proteins. If another marker is necessary, the HaloTag of the antiGFP-nanobody-HaloTag-iLID construct can be left unstained. Additionally, a nanobody against RFP is available, making the system adjustable to RFP-tagged target proteins as well (Prole and Taylor, 2019).

If there is a cell line available of the POI tagged with GFP or RFP expressed on the endogenous level, the antiGFP-nanobody-HaloTag-iLID construct could be transiently transfected and can bind the GFP-tagged protein, without the necessity to overexpress it or create a new stable cell line. However, the active signaling protein is often fused to the SspB. To avoid overexpression of signaling molecules, one can use the same principle but create a SspB-antiGFP-nanobody construct and use an endogenously GFP tagged signaling protein to be bound by an antiGFP-nanobody-SspB.

In this study, we show the effective relocalization of SspB induced by light to iLID localized at the nucleus, ER and plasma membrane via the antiGFP nanobody. Previous work by Küey and co-workers showed the efficient relocation of an antiGFP nanobody to a GFP-tagged protein, here a mitochondrial tag, in combination with the rapamycin heterodimerization system (Küey et al., 2019). However, the same study finds that the binding of the nanobody to dynamin-2-GFP, without the activation of the rapamycin system, inhibits the clathrin-mediated endocytosis. Thus, the alteration of protein function through nanobody binding is a potential limitation and it is necessary to study the potential side-effects before applying the nanobody as a linker. Nevertheless, a study by Hansen and colleagues found that the direct fusion of the optogenetic tool resulted in a loss of function and the nanobody-based targeting enabled the proper localization in the primary cilium (Hansen et al., 2020).

One other limitation to take in account, is the potential accumulation of unbound nanobody in the cytosol. This would occur if the nanobody construct is expressed at a higher level than the GPF construct. Subsequently, all the available GFPs are bound by the nanobody, so the superfluous molecules would accumulate in the cytosol and lead to untargeted SspB recruitment during activation. One solution is to adjust the expression level of the nanobody construct through a different promoter. However, a more elegant solution has been presented by Oliinyk and colleagues in form of an antigen-dependent nanobody (Oliinyk et al. 2022). They inserted the fluorescent tag, miRFPnano3, in a nanobody against the protein of interest. In the unbound form this construct is ubiquitinated and degraded, resulting in the removal of unbound probe. Binding of the nanobody to its epitope stabilizes the construct. Another limitation might be the size and bulkiness of the GFP bound to the Nanobody(antiGFP)-HaloTag-iLID construct fused to the protein of interest, which may compromise its functionality and localization. This issue could be tackled by using the smaller miRFP670nano3 near-infrared tag instead of the larger HaloTag (Oliinyk et al. 2022).

Aiming toward the optogenetic manipulation of endogenous proteins, the nanobody-targeting strategy offers different options. If a nanobody exists for the protein of interest, a nanobody(antiPOI)-SspB could be applied together with iLID target to the desired location in the cell. This strategy would allow the recruitment of the endogenous protein. However, the effects of the nanobody binding should be carefully tested in comparison to the effects of the recruitment. Any manipulation of the system will require some alterations, which may have side-effects. Another option is the tagging of the POI in its location in the genome with the iLID system but in this case the entire fusion protein has to be transcribed and potentially post transcriptional modified, whereas the nanobody would bind the already expressed protein potentially interfering less with its maturation.

Recently, light-switchable nanobodies (OptoNBs) have been described (Gil et al., 2020). These nanobodies bind or unbind their target when exposed to blue light and they are based on an engineered LOV domain. If such an OptoNB could be applied for the POI, it would allow the undisturbed maturation of the protein and only at the moment of the light exposure, the OptoNB would bind and interfere with the endogenous situation preventing adaptation of the cell or basal activity of an overexpressed component. This OptoNB could be localized directly at the desired location and recruit the POI to it. However, the amount of specific nanobodies is still limited especially for OptoNBs. There are OptoNBs against GFP and mCherry. It has been demonstrated that OptoNB is able to recruit the ERK regulator SOS(cat) to the plasma membrane tagged mCherry (Gil et al., 2020). In that case, all components were overexpressed but in principle, this method could be used to recruit endogenously fluorescent protein-tagged proteins and the combination with the iLID system might increase recruitment efficiency.

Optogenetics and nanobodies are a powerful combination that could enable spatial and temporal precise manipulation of endogenous proteins. The hybrid developed in this study can be used to recruit a protein of interest to a GFP fusion protein and represents just one out of many options. We anticipate the engineering of more combinations, further increasing the options and improving flexibility and control of optogenetic manipulations.

## Methods

### Plasmids

#### Nanobody

The plasmid containing the anti-GFP nanobody was a gift from Brett Collins (addgene plasmid # 49172). nanobody(antiGFP)-mScarlet-I-N1 was created by PCR amplifying the insert nanobody(antiGFP) with the primers: FW-5’TATAGGTACCATGCAGGTTCAACTGGTG3’ and RV-5’ TATAACCGGTGAAGAGCTCACCGTCACCTG3’, digesting it with KpnI and AgeI and ligating it to the likewise digested backbone mScarlet-I-N1. mScarlet-I-nanobody(antiGFP) was created by PCR amplifying the insert nanobody(antiGFP) with the primers: FW-5’TATATGTACAAGTCCATGCAGGTTCAACTGGTG3’ and RV-5’TATAGGTACCTTAAGAGCTCACCGTCACCTG3’, digesting it with BsrGI and KpnI and ligating it to the likewise digested backbone mScarlet-I-C1. mNeonGreen-nanobody(antiGFP) was created by digesting mNeonGreen with AgeI and BsrGI and ligating it to the likewise digested backbone mScarlet-I-nanobody(antiGFP).

#### H2A

The plasmid LSS-sGFP2-H2A-C1 is available on addgene (plasmid #112937). iRFP670-H2A, EGFP-H2A, mNeonGreen-H2A, mTurquoise2-H2A, mScarlet-I-H2A, mKate2-H2A, mCherry-H2A were created by digesting the inserts: iRFP670, mNeonGreen, mTruquoise2, mScarlet-I, mKate2, and mCherry with AgeI and BsrGI and ligating them to the likewise digested backbone LSS-sGFP2-H2A-C1.

#### iLID

The cysteine-free, or secretory HaloTag version 7 was provided by Ariana Tkachuk in consultation with Erik Snapp (Janelia Research Campus of the Howard Hughes Medical Institute). The HaloTag was PCR amplified (primers: FW 5’-ATATACCGGTCGCCACCATGGCCGAGATCGGCA-3’ and RV 5’-ATATTGTACACGCCGCTGATCTCCAGG-3’) and AgeI, BsrGI digested and ligated into a likewise digested C1 (Clontech) backbone, creating C1-HaloTag. C1-Lck-mTurquoise2-iLID was described previously and is available on addgene (plasmid #176125). Nanobody(antiGFP)-HaloTag-iLID was created by first digesting the backbone Lck-mTurquoise2-iLID with AgeI and BsrGI and ligating it to the likewise digested insert HaloTag, thereby creating Lck-HaloTag-iLID. To subsequently create Nanobody(antiGFP)-HaloTag-iLID, the backbone Lck-HaloTag-iLID was digested with AgeI and NdeI and ligated to the likewise digested insert nanobody(antiGFP) from nanobody(antiGFP)-mScarlet-I-N1. SspB-mScarlet-I-C1 was described previously (Van Geel et al., 2018). The, in this study, created plasmid Nanobody(antiGFP)-HaloTag-iLID (addgene# 191453) is available on addgene.

#### Other

The plasmid containing CytERM-mScarlet is available on addgene (plasmid # 85066). To create CytERM-mEGFP, mEGPF was digested with AgeI and BsrGI and ligated to the likewise digested backbone CytERM-mScarlet-N1. EGFP-CaaX-C1 is available on addgene (plasmid # 86056).

### Cell Culture and Transfection

Henrietta Lacks (HeLa) cells (CCL-2, American Tissue Culture Collection) were cultured in Dulbecco’s modified Eagle’s medium + GlutaMAX™ (Giboc) with 10% fetal calf serum (Giboc) (DMEM + FCS) at 37°C and 7% CO_2_. For transfection 25 000 to 50 000 HeLa cells per well were seeded on round 24 mm ø glass coverslips (Menzel, Thermo Fisher Scientific) in a 6 well plate with 2 ml DMEM + FCS or alternatively roughly 10 000 HeLa cells per well were seeded in a glass bottom 24-well No 1.5, uncoated (MatTek Corporation) in 1 ml DMEM + FCS. The Transfection mix contained 1μl linear polyethylenimine (Polysciences) per 100 ng DNA, at a concentration of 1 mg/ml and a pH of 7.3, and for the 6-well plate the mix contained 0.5 μg plasmid DNA, mixed with 200 μl OptiMEM (Thermo Fisher Scientific) per well, for the 24-well plate it contained 125 ng plasmid DNA and 50 μl OptiMEM per well. After 15 min incubation at room temperature the transfection mix was added to the cells, 24 h after seeding.

### Confocal Microscopy

The HaloTag was stained, for at least 1 h prior to imaging, with a concentration of 150 nM of Janelia Fluor Dyes or JF635 nm (far red) (Janelia Materials) in the culture medium. HeLa cells were imaged between 16 and 24 h after transfection either directly in the glass bottom 24-well plate, or on the 24 mm ø glass coverslips transferred to an Attofluor cell chamber (Thermo Fischer Scientific), in 1 ml of Microscopy Medium (20 mM HEPES (pH=7.4),137 mM NaCl, 5.4 mM KCl, 1.8 mM CaCl_2_, 0.8 mM MgCl_2_ and 20 mM Glucose) at 37°C. Confocal microscopy images were obtained at a Leica Sp8 (Leica Microsystems) using unidirectional line scan at a scan speed of 400 Hz. Images were acquired with 1024×1024 pixel resolution and 16-bit color depth.

H2A construct anti-GFP nanobody colocalization was imaged with a 20x objective (HC PL Apo, CS2, NA=0.75, dry), pinhole set to 2 Airy Unit, 8 x line averaging and the fluorescent signal was detected sequentially. mTurquoise2 signal was measured with a 442 nm diode laser line in combination with at HyD detector, emission detection range set to 452 – 552 nm and gain set to 110 V. LSS-sGFP2 signal was measured with a 442 nm diode laser line in combination with a HyD detector, emission detection range set to 498 – 552 nm and gain set to 100 V. mNeonGreen and EGFP signal was measured with a 488 nm argon laser line in combination with a HyD detector, emission detection range set to 498 – 552 nm and gain set to 100 V. mCherry and mKate2 signal was measured with a 594 nm HeNe laser line in combination with a HyD detector, emission detection range set to 604 −704 nm) and gain set to 50 V. mScarlet-I signal was measured with a 561 nm DPSS laser line in combination with a HyD detector, emission detection range set to 571 −671 nm, gain set to 50 V. iRFP670 signal was measured with a 633 nm HeNe laser line in combination with a HyD detector, emission detection range set to 644 −744 nm, gain set to 100 V.

Optogenetic recruitment was imaged with a 63x objective (HC PL Apo, C2S, NA=1.40, oil), pinhole set to 1 Airy Unit, 2x line averaging and a frame interval of 10 s. Photo activation was achieved with a 488 nm argon laser set to 20% laser power and 1% intensity with 2x line averaging scanning the whole field of view every 10 s for 4 min. The EGFP signal was only measured during photo activation. Far red HaloTag JF635 nm and EGFP signal were measured simultaneously using a 488 nm argon laser line and a 633 HeNe laser line in combination with a PMT detector with an emission detection range set to 498 to 550 nm and a gain of 600 V and a HyD detector with an emission detection range of 643 – 743 nm and the gain set to 100 V. mScarlet-I signal was measured sequentially using a 561 nm DPSS laser line in combination with a HyD detector set to an emission range of 571 – 630 nm with a gain of 100 V.

### Analysis

FIJI was used to prepare representative microscopy images by adjusting brightness and contrast (Schindelin et al., 2012).

The nuclear to cytosolic intensity ratio was measured with CellProfiler (Version 4.1.3) (Stirling et al., 2021). The pipeline can be found on in the data deposit on Zenodo. Raw images were uploaded to CellProfiler. The primary object, representing the ROI of the nucleus, was identified in the H2A-fluorescent protein images for objects between 30 and 100 pixel diameter with the Otsu thresholding method. If necessary, the identified ROIs of the nucleus were corrected manually. The ROI for the cytosol was defined by extending the ROI of the nucleus by 1 pixel and then subtracting the ROI of the nucleus. The intensity was measured for the ROIs of the nucleus and the cytosol and the ratio of nuclear over cytosolic intensity was calculated. The ratio was plotted with the web-tool ‘PlotsofData’ and the 95% confidence intervals were calculated by bootstrapping (Postma and Goedhart, 2019).

For the membrane recruitment ROIs for the plasma membrane and the cytosol were drawn by hand in a region with little cell movement. For nuclear recruitment ROIs for the nucleus and the cytosol were drawn by hand. For ER recruitment, the image was cropped so that entire field of view was covered by cell, a Gaussian blur (sigma radius=0.5) filter was applied, the ‘MinError’ thresholding method was applied to define the ROI of the ER for each frame. This ROI was inverted to represent the cytosol. SspB-mScarlet-I intensity for nucleus, plasma membrane, ER and cytosol ROIs was measured. The ratio of either nuclear, plasma membrane or ER intensity over cytosolic intensity was calculated and plotted over time in the web-tool ‘PlotTwist’ (Goedhart, 2020).

## Acknowledgements

We want to thank Ronald Breedijk for the support at the van Leeuwenhoek Centre for Advanced Microscopy, Section Molecular Cytology, Swammerdam Institute for Life Sciences, University of Amsterdam.

## Author contribution

E.M. conceptualized, investigated, analyzed, visualized, and wrote the manuscript, M.T. and T.H. provided resources; J.G. acquired funding, supervised, conceptualized the project and reviewed and edited the manuscript.

## Data availability

The data generated during this study is available at Zenodo.org (after publication): DOI: 10.5281/zenodo.7152304

